# Individual variability in functional organization of the human and monkey auditory cortex

**DOI:** 10.1101/2020.01.06.895474

**Authors:** Jianxun Ren, Hesheng Liu, Ting Xu, Danhong Wang, Meiling Li, Yuanxiang Lin, Julian S.B. Ramirez, Jie Lu, Luming Li, Jyrki Ahveninen

**Affiliations:** Athinoula A. Martinos Center for Biomedical Imaging, Department of Radiology, Massachusetts General Hospital, Harvard Medical School, Charlestown, MA, USA; National Engineering Laboratory for Neuromodulation, School of Aerospace Engineering, Tsinghua University, Beijing, China; Department of Neuroscience, Medical University of South Carolina, Charleston, SC, USA; Center for the Developing Brain, Child Mind Institute, New York, NY, USA; Department of Neurosurgery, First Affiliated Hospital, Fujian Medical University, Fuzhou, China; Department of Behavior Neuroscience, Department of Psychiatry, Advanced Imaging Research Center, Oregon Health and Science University, Portland, OR, USA; Department of Radiology, Xuanwu Hospital, Capital Medical University, Beijing, China; Precision Medicine & Healthcare Research Center, Tsinghua-Berkeley Shenzhen Institute, Tsinghua University, Shenzhen, China; IDG/McGovern Institute for Brain Research at Tsinghua University, Beijing, China

**Author notes:** Address correspondence to: Jie Lu, M.D., Ph.D., Department of Radiology, Xuanwu Hospital, Capital Medical University, Beijing, China, Or Luming Li, Ph.D., National Engineering Laboratory for Neuromodulation, School of Aerospace Engineering, Tsinghua University, Beijing, China. Co-first Authors (Jianxun Ren, Hesheng Liu).

**Keywords:** auditory cortex, individual differences, nonhuman primate, functional connectivity

## Abstract

Accumulating evidence shows that auditory cortex (AC) of humans, and other primates, is involved in more complex cognitive processes than feature segregation only, which are shaped by experience-dependent plasticity and thus likely show substantial individual variability. However, thus far, individual variability of ACs has been considered a methodological impediment rather than a phenomenon of theoretical importance. Here, we examined the variability of ACs using intrinsic functional connectivity patterns in humans and macaques. Our results demonstrate that in humans, functional variability is 1) greater near the non-primary than primary ACs, 2) greater in ACs than comparable visual areas, and 3) greater in the left than right ACs. Remarkably similar modality differences and lateralization of variability were observed in macaques. These connectivity-based findings are consistent with a confirmatory task-based fMRI analysis. The quantitative proof of the exceptional variability of ACs has implications for understanding the evolution of advanced auditory functions in humans.

## Introduction

Association areas of brain that underlie complex cognitive qualities such as speech and language demonstrate considerable individual variability (Mueller et al., 2013; Stoecklein et al., 2019). In contrast, sensory areas of the cerebral cortex, which are evolutionarily old (Kaas, 2006) and maturate at early stages of human development (Hill et al., 2010), have been considered to be relatively similar across individuals. Emerging evidence, however, suggests that the auditory system represents an exception to this rule (King and Nelken, 2009). The basic attributes of auditory stimuli are processed much more thoroughly in subcortical nuclei than those of visual stimuli (Masterton, 1992). Even in primary ACs, neurons have dense integrative lateral connections (Lu and Wang, 2004) and strong preference for complex sound patterns rather than isolated features only (Moerel et al., 2013; Nelken, 2004). In contrast to early visual cortex (VC) areas, relatively early aspects of ACs are involved in complex perceptual functions, such as speech and music processing (Griffiths and Warren, 2002; Mesgarani et al., 2008; Norman-Haignere et al., 2015), which are strongly modified by the individuality of our lifelong experiences (Herholz and Zatorre, 2012; Ressel et al., 2012). Systematic investigation of individual functional variability could, thus, offer a way to examine the hierarchical arrangement of human ACs and to enhance our understanding of how their unique properties differ from other sensory areas of the brain.

Previous studies have considered individual variability of ACs as a methodological impediment rather than a phenomenon of theoretical importance. Pioneering studies of human AC anatomy, which were based on three-dimensional (3D) stereotactic anatomical normalization, were complicated by the substantial individual variability of Heschl’s gyrus (HG), the primary anatomical landmark of ACs (for a review, see Moerel et al., 2014). Today, this problem can be greatly alleviated thanks to more precise surface-based inter-subject alignment methods (Coalson et al., 2018; Fischl and Sereno, 2018), as recently verified by using non-invasive measures of AC myeloarchitecture (De Martino et al., 2015; Dick et al., 2012). Functional alignment of AC areas has, in turn, remained a problem due to the lack of a definite localizer paradigm. Whereas the subarea boundaries of VC can be functionally mapped based mirror-symmetric representations of the visual field polar angle and eccentricity (Sereno et al., 1995), in AC the problem is that the topographic representation of cochlea is one dimensional. Although great advances in our understanding of human AC have been recently achieved by using novel data-driven approaches (Kell and McDermott, 2019; Moerel et al., 2013; Norman-Haignere et al., 2015), the exact layout of AC still remains an open question. Due to the lack of unequivocal mapping paradigm, the degree of individual variability of different AC areas has also remained a widely shared belief rather than quantified fact.

A powerful way to characterize the individuality of our brains, which has so far been largely unexploited in human AC mapping, is the analysis of their functional connectome (Seung, 2012). In previous studies, such analyses have been conducted using resting state functional connectivity MRI (fcMRI) (Mueller et al., 2013; Stoecklein et al., 2019). A remarkable and highly replicable finding of these fcMRI studies has been that despite their variability at the group level, within any individual brain the intrinsic functional connectivity patterns are highly robust and reliable, to a degree that a specific person can be identified from a larger group of subjects based on fcMRI (Finn et al., 2015). The smaller number of fcMRI studies that have so far been conducted in the auditory domain show that, consistently with neurophysiological recordings in VCs (Kenet et al., 2003), within the early ACs the intrinsic functional connectivity patterns are consistent with feature-topographic pathways (Cha et al., 2016; Lumaca et al., 2019). The inter-individual variability, and the within-individual diversity of longer-range connections across neighboring voxels, could however significantly increase as a function of hierarchical level (Mueller et al., 2013). Thus, by controlling for the variability of anatomical properties such as cortical folding measures, as well as for the noise introduced by within subject functional variability, it could possible to estimate the individual variability of different levels of AC processing and to compare it to other sensory areas, independent of anatomical biases and regional differences in MRI data quality (Mueller et al., 2013).

The inter-individual differences in AC could be related to understanding of the evolution of our unique, human-specific auditory-cognitive skills. There is increasing evidence that not only humans, but also non-human primates show communication behaviors that cannot be explained without the existence of a highly advanced auditory system (Belin, 2006; Ghazanfar and Santos, 2003). For example, the vocalizations that non-human primates use for group communication show subtle but rich variability depending on the social context (Aboitiz, 2018), across different populations of the same subspecies (Arcadi, 1996), and even between different individuals within a specific population (Salmi et al., 2014). The ability to interpret these modulations has evolved alongside an increasingly complex ACs (Hackett et al., 2001), which has a strong capacity for adaptive plasticity (Cheung et al., 2005) and which, thus, also likely show considerable functional variability between individuals.

Here, to elucidate the individual variability in the functional organization of the AC, we quantified variability based on resting state connectivity patterns and investigated whether variability increases as a function of processing hierarchy in individual subjects. In all these analyses, we used the individual variability of cortical folding patterns as well as the within-subject variability of functional connectivity estimates as covariates, to control for biases caused by regional differences in anatomical variability, physiological noise, and MRI data quality. We further tested whether inter-subject variability is greater in the AC than in VC and reflects some features of higher-order processing, such as hemispheric lateralization. The results obtained in humans were compared to a resting-state fMRI data obtained in the macaque, which offered a way to verify the inter-species consistency of AC vs. VC differences in a model that lacks the additional 3D variability caused by HG, a structure that is found only in humans.

## Results

### Substantial inter-subject variability in the human and macaque auditory cortex

Functional connectivity, and its individual variability, was estimated in human AC using a resting-state fMRI dataset that consists of 30 young healthy adults (the CoRR-HNU dataset (Zuo et al., 2014), 15 females, age 24 ± 2.41 yrs). Each subject underwent ten scanning sessions (10 min resting-state fMRI each session, i.e., 100 min fMRI data per subject, see Materials and Methods) over approximately one month. For each vertex in the AC, its connectivity with all other vertices in the cerebral cortex was calculated using the data of each session and then averaged across 10 sessions. Inter-subject variability of functional connectivity was quantified at each vertex based on the dissimilarity of the seed-based connectivity maps between subjects, using the strategy described in (Mueller et al., 2013). Specifically, to control for the impact of noise and other technical confounds, inter-subject variability in connectivity was corrected by linearly regressing out the mean intra-subject variability (Mueller et al., 2013), which was quantified in each subject based on the variation of connectivity maps across 10 sessions. We replicated the previous finding of inter-subject variability in functional connectivity in the human brain, which indicated high variability in the association cortices but low variability in the visual and sensorimotor areas (Figure 1A). Values below the global mean are shown in cool colors, while values above the global mean are shown in warm colors.

**Figure 1.**
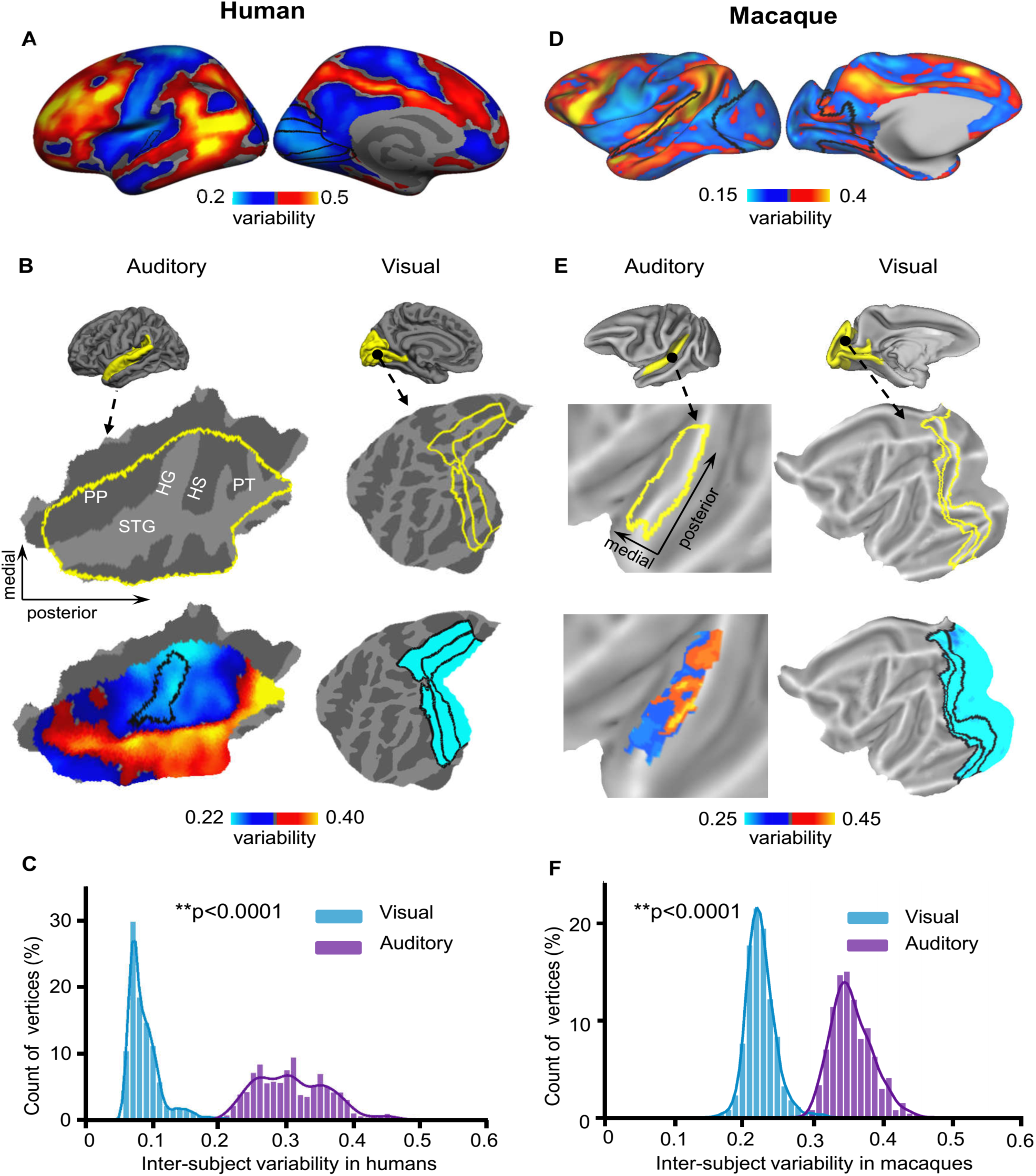
Inter-individual variability in the AC is significantly larger than that in the visual cortex in both humans and macaques. **(A)** Inter-individual variability in functional connectivity derived from the CoRR-HNU dataset (N=30) is shown in the human cortical surface. **(B)** The auditory cortex (AC, left column) and the visual cortex (VC, right column) are displayed as magnified flattened patches. Inter-individual variability in AC and VC is plotted. Variability is much larger in AC than in VC. HG: Heschl’s gyrus; PT: planum temporale; STG: superior temporal gyrus; PP: planum polare. **(C)** Histograms of inter-subject variability in the AC (purple bars) and VC (blue bars). The AC shows significantly higher inter-subject variability than the VC (p < 0.0001, Wilcoxon Rank Sum test). **(D)** Inter-individual variability in functional connectivity derived from the Macaque dataset (N=4) is shown in the macaque cortical surface. Variability in macaques demonstrates the similar principle of the spatial distribution with that in humans. **(E)** Inter-individual variability is shown in the macaque AC and VC. In macaques, AC also demonstrates much greater variability than VC. **(F)** Histograms of variability show that AC (purple bars) more variable than VC (blue bars, p < 0.0001, Wilcoxon Rank Sum test) in macaques.

Focusing on the auditory cortex, we found that inter-subject variability is relatively low in Heschl’s gyrus (HG) but much greater laterally in the superior temporal gyrus (STG), which could be near the human homolog of monkey parabelt areas (Figure 1B). This suggests that the non-primary auditory areas may be more variable across individuals than the primary auditory areas. Seed-based connectivity analysis indicated that a region in the low variability area is strongly connected to the sensorimotor cortex, whereas a nearby region in the high variability area shows strong connectivity to the frontal lobe (see Figure S1). For comparison purposes, we then quantified inter-subject variability in the VC (Figure 1B). Critically, we found that inter-subject variability in the AC is significantly larger than that in the VC (Figure 1C, p<0.0001, Wilcoxon Rank Sum test, the curves represent fitted data using a kernel distribution). We then replicated the findings in an independent dataset (MSC dataset) (Gordon et al., 2017), which included 10 healthy young adults (5 females, age 29.1 ± 3.3 yrs). Each subject underwent 10 scanning sessions (30 min resting state fMRI each session, see Materials and Methods) on 10 separate days. Although the two datasets differ in the subjects’ ethnicities and scanning parameters, we found that the spatial distribution of inter-subject variability in the AC was highly replicable (r = 0.836, p <0.0001, see Figure S2A & S2B).

We then investigated inter-subject variability in functional connectivity across four macaque monkeys. Two subjects were scanned for eight 10-min fMRI runs under anesthesia (see Materials and Methods) and the other two subjects were scanned for eight 30-min fMRI runs under anesthesia (Xu et al., 2018) but only the first 10-min of each run was retained for analyses thus the data length was kept the same for all subjects. The procedure for evaluating inter-subject variability in macaque is identical to the procedure for the human data as described above (see Materials and Methods and Figure S3 for the definition of auditory mask in macaques (Markov et al., 2012)). We found that inter-individual variability in macaque monkeys demonstrated the similar principal of the spatial distribution with that in humans, i.e., associated areas in the frontal, parietal and temporal lobes show marked inter-individual variability while primary areas such as sensorimotor and VCs demonstrate low variability. Note that the color scale of variability has been scaled differently for two species so the gradient within each species can be better appreciated (Figure 1D). Importantly, the macaque auditory areas showed substantial inter-subject variability (Figure 1E), which is significantly higher than that in the VC (Figure 1F, p<0.0001, Wilcoxon Rank Sum test).

### Lateralization of inter-individual variability in the AC

One of the important functions of the human AC is speech processing, which is lateralized at the population level but varies across individuals. Here, we investigated whether ACs in two hemispheres show similar levels of individual variability, or if one hemisphere is more variable than the other. Inter-subject variability in functional connectivity was quantified in the left and right ACs using the CoRR-HNU dataset (Figure 2A). While both hemispheres showed similar spatial distributions of inter-subject variability with low variability in HG and high variability near the STG, variability is much greater (p<0.001, Wilcoxon Rank Sum test) in the left AC than in the right AC (Figure 2B). This finding was then successfully replicated in the MSC dataset (p<0.001, Wilcoxon Rank Sum test, see Figure S2c & S2D). These observations imply that the left AC may be more involved in higher-order functional processing than the right AC.

**Figure 2.**
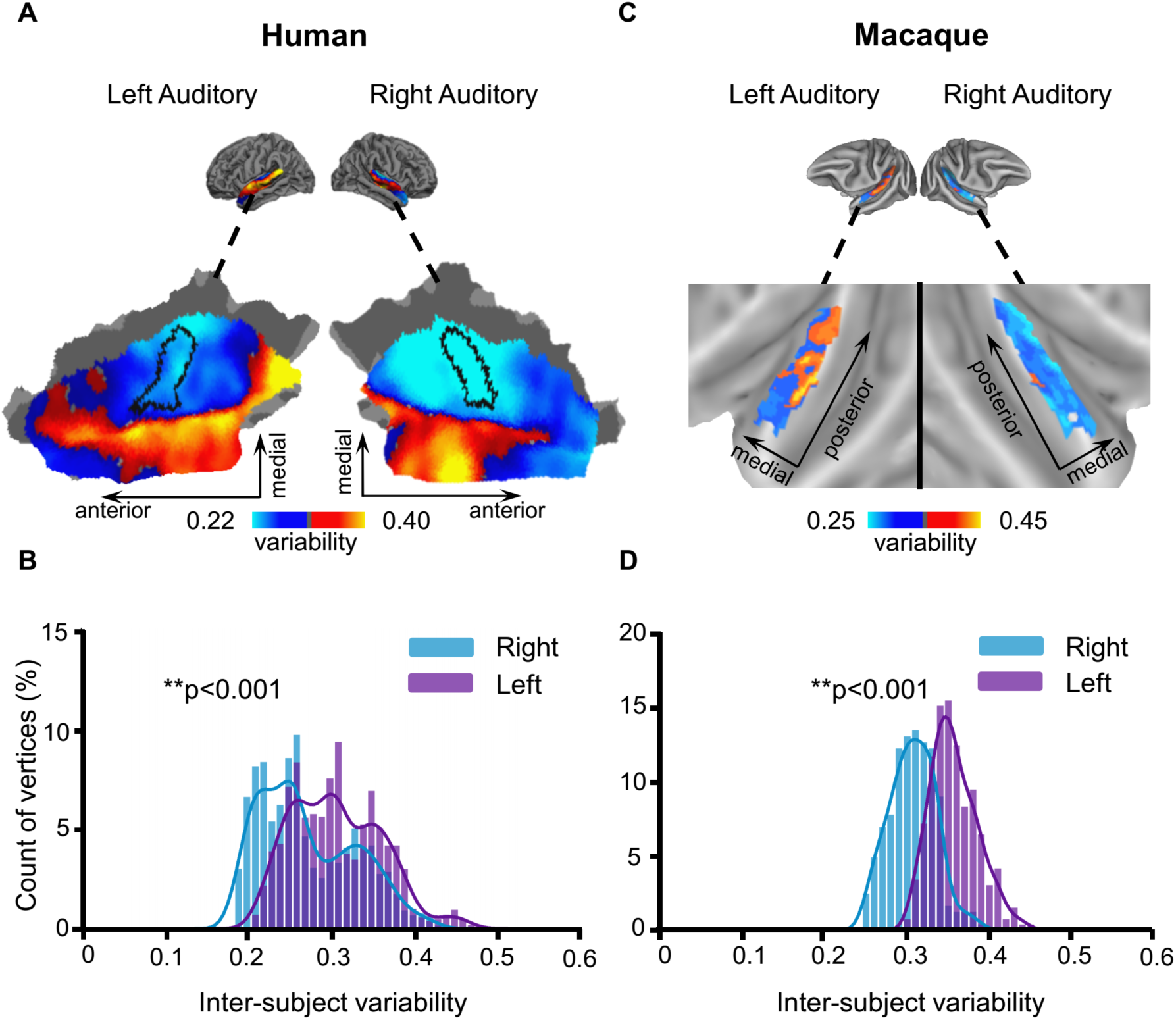
Inter-individual variability in the AC shows significant lateralization in both humans and macaques, with the left AC being more variable than the right. **(A)** Spatial distribution of inter-subject variability in the left and right ACs of humans. Values below the mean within both left and right AC are shown in cool colors, while values above the global mean are shown in warm colors. Variability appears to be higher in the left AC than in the right AC. **(B)** Histograms of inter-subject variability in the left and right ACs of humans show significantly higher inter-individual variability in the left AC (purple bars) than in the right AC (blue bars, p < 0.001, Wilcoxon Rank Sum test). **(C)** Spatial distribution of variability in the left and right ACs of macaques **(D)** Histograms of inter-subject variability in the left and right ACs of macaques also show significant left lateralization (p < 0.001, Wilcoxon Rank Sum test).

We next investigated whether inter-individual variability is lateralized in the macaque AC. Strikingly, in macaques, left AC also demonstrated significantly greater variability than right AC (Figure 2C & 2D, p<0.001, Wilcoxon Rank Sum test), indicating the lateralization pattern observed in the human AC might have an evolutionary trace.

### Inter-subject variability in task-evoked activations in the human AC

Recent studies have indicated that individual differences in resting state connectivity are related to individual differences in task-evoked activity (Tavor et al., 2016). Here, we examined whether the spatial distribution of individual variability in the AC could also be observed in task-evoked activity. Inter-subject variability in task-evoked fMRI activations was assessed in the AC using the *Human-voice dataset* (N=218, see Materials and Methods). Subjects were scanned while passively listening to vocal and non-vocal stimuli (Pernet et al., 2015). Inter-subject variability was estimated as the standard deviation of the *z*-values from task activation across all subjects, with the mean *z*-values regressed out (see Figure S4). Interestingly, we also found low inter-subject variability in HG and higher variability in the lateral STG (i.e., the possible human homolog of the monkey parabelt area). Furthermore, task fMRI variability of both vocal and non-vocal stimulus were significantly correlated with resting-state functional connectivity variability (Figure 3A & Figure 3C, for non-vocal stimulus, r = 0.504, p <0.0001; for vocal stimulus, r = 0.502, p < 0.0001). Moreover, inter-individual variability in task-evoked fMRI activations in ACs also showed left lateralization. Left AC demonstrated significantly greater variability in task-evoked activity than right AC (Figure 3B and Figure 3D, for both stimuli, p < 0.001, Wilcoxon Rank Sum test).

**Figure 3.**
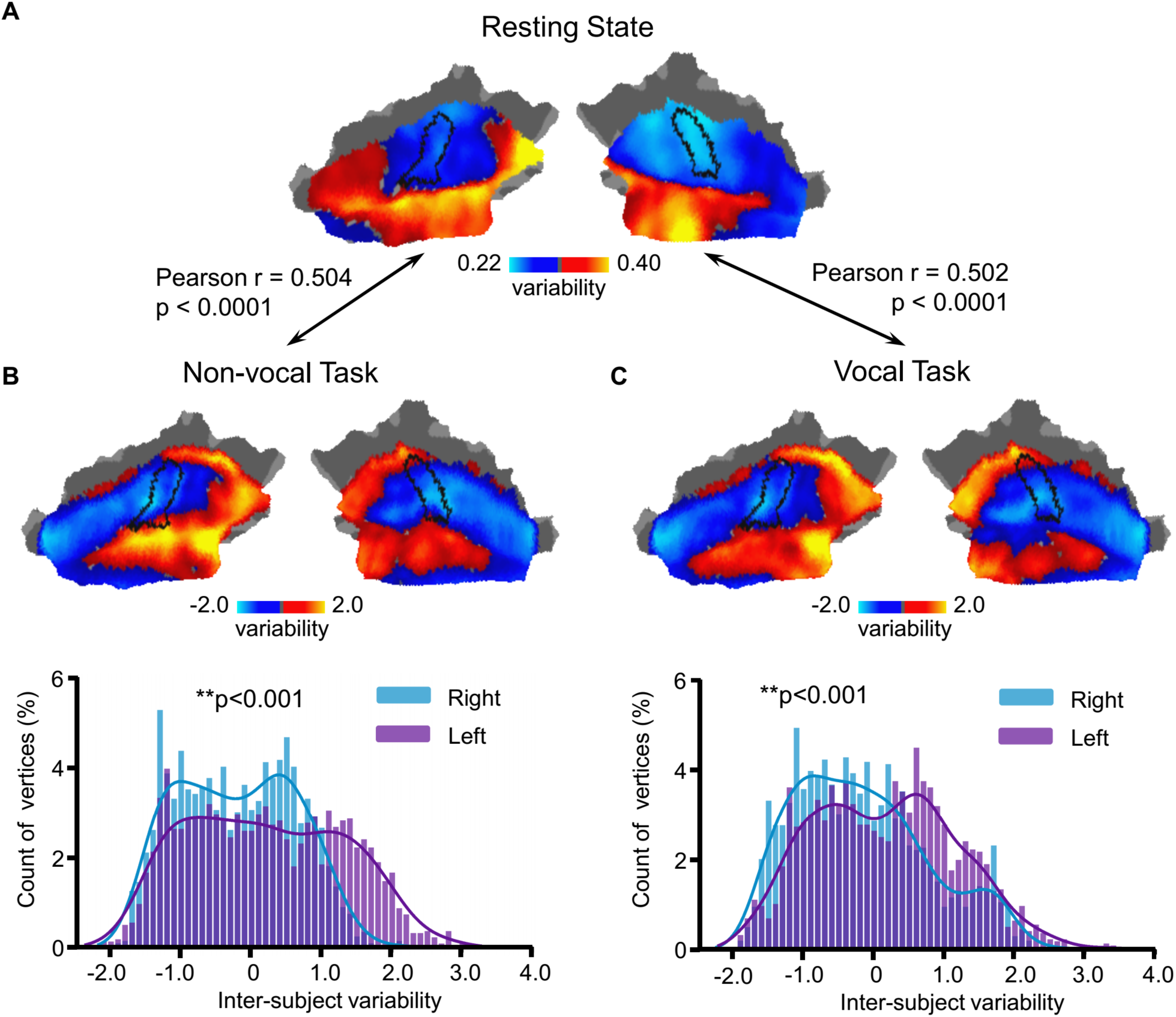
Inter-individual variability in task-evoked activations in the AC shows the same principle of the spatial distribution as individual variability in functional connectivity. Inter-subject variability in task-evoked fMRI activations (*Human-voice dataset*, N = 218) was assessed in the AC. Variability was estimated as the standard deviation of the *z*-values from task activation across all subjects, with the average *z*-values regressed out. Inter-individual variability estimated at rest **(A)** and variability estimated using task activations based on non-vocal **(B)** and vocal **(C)** auditory stimuli, show the same principle of the spatial distribution, with low inter-subject variability near Heschl’s gyrus (indicated by a black curve) but higher variability in the lateral superior temporal cortex (i.e., the likely human parabelt area). Variability derived from task activations is correlated with the variability estimated at rest (r = 0.504, p <0.0001 for non-vocal auditory stimuli and r = 0.502, p < 0.0001 for vocal auditory stimuli). Moreover, inter-individual variability in task-evoked activations in the ACs also show significant left lateralization. The histograms of variability estimated using both tasks **(B, C)** indicate that the left AC (purple bars) show significantly higher variability than the right AC (blue bars, p < 0.001, Wilcoxon Rank Sum test).

### Relationship between functional and anatomical variability in the AC

Anatomical variability in the human AC has been well recognized in the literature. We therefore investigated how functional variability may be related to known anatomical variability. Inter-subject variability in sulcal depth and cortical thickness was assessed using intraclass correlation (ICC), with intra-subject variance properly accounted for (Mueller et al., 2013). We found that variability in functional connectivity showed a moderate correlation with variability in sulcal depth (Figure 4, r = 0.36, p <0.0001), but not with cortical thickness (r = −0.04, p = 0.084).

**Figure 4.**
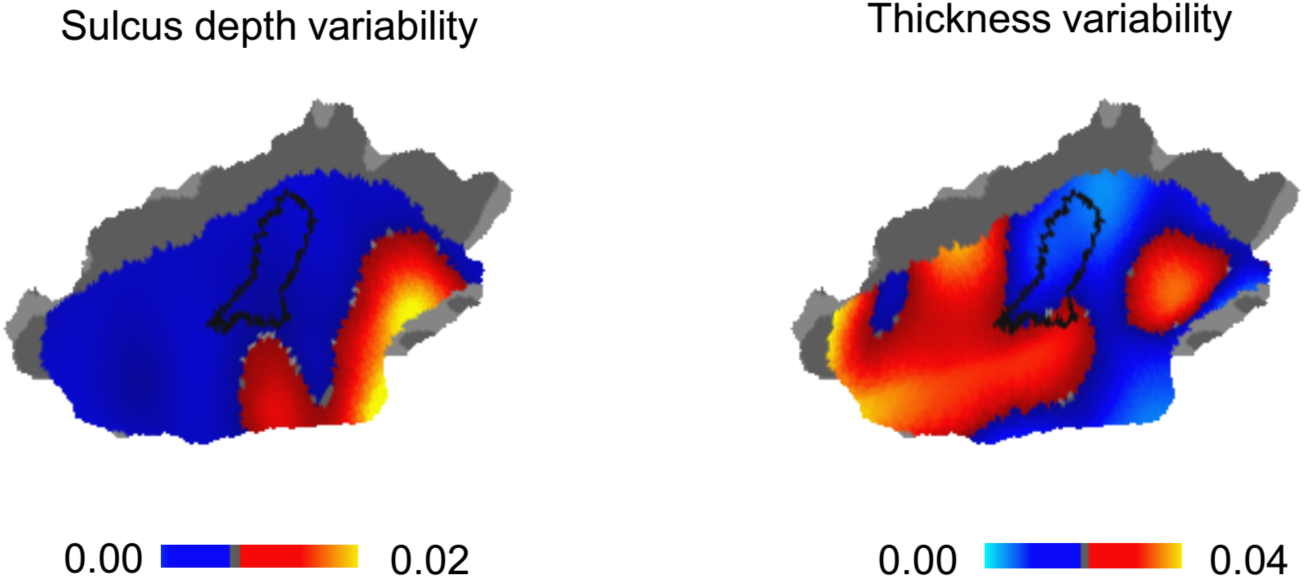
Functional Variability in the human AC is moderately associated with variability in sulcal depth, but not cortical thickness. Inter-individual variability in sulcal depth (left) and cortical thickness (right) was assessed using intraclass correlation (ICC), with intra-individual variance properly accounted for. Variability in functional connectivity is moderately associated with variability in sulcal depth (Pearson correlation r = 0.36, p <0.0001), but not with cortical thickness (Pearson correlation r = −0.04, p = 0.084).

## Discussion

Unveiling the complex functional organization in the human AC remains a major challenge in neuroscience research, largely due to marked individual variability in the AC. Here, we used resting state and task-based fMRI to investigate the individual variability of AC functions in humans and macaque monkeys. The results reveal a unique spatial distribution of variability in the AC that likely follows the auditory processing hierarchy, i.e., regions near the primary auditory areas demonstrated lower individual variability than non-primary areas. Compared to the VC, the AC demonstrated much greater individual variability in functional connectivity, suggesting that certain parts of the AC are more similar to higher-order association areas than early sensory regions in both primate species. Furthermore, we found that the left AC is more variable than the right AC, which may be related to its role in some lateralized, higher-order functions, such as primate auditory-vocal communication that has evolved to speech and language in humans. The spatial distribution of individual variability in AC function could also be observed using task fMRI data in humans, confirming that non-primary AC areas are more variable than primary AC areas. Taken together, our findings reveal a putative functional hierarchy in the primate AC and indicate that portions of the AC are particularly variable across individuals and possess some characteristics of areas associated with complex cognitive functions.

### Functional hierarchy in the human and macaque AC

One of the most widely accepted organizational principles of sensory systems is parallel/hierarchical processing. Different stimulus attributes are first segregated to separate pathways, and then integrated step by step to increasingly complex object representations. Our previous work demonstrates that functional connectivity is relatively consistent across individuals in those unimodal sensory and visual areas but varies substantially in multimodal association areas (Mueller et al., 2013). Inter-subject variability is also closely related to evolutionary expansion, developmental expansion, and hemispheric specialization (Wang et al., 2015). The gradient of inter-subject variability in the primate brain may thus reflect the hierarchy of functional processing. Focusing on the human AC, our present results demonstrate in two independent data sets that functional connectivity is significantly more variable in the lateral part of the AC in the STG than in areas close to the medial HG (Figure 1 and Figure S2). This sharp transition of variability was also observed in task fMRI data (Figure 4). These observations suggest that functional complexity may abruptly increase near the STG. More detailed future studies using similar methods could thus provide critical information about the functional hierarchy in the human AC, which has so far been much more difficult to specify than that in the visual and somatosensory cortices, resulting in differing interpretations of how the human AC processes information (Bizley and Cohen, 2013), including spoken language (for a review, see Rauschecker and Scott, 2009).

Previous studies have also provided evidence that a consistent pattern of functional mapping results is harder to replicate across different subjects in non-primary than primary AC areas (Moerel et al., 2014). However, in many of these previous studies, there were a number of alternative explanations that could have accounted for the increased inter-subject variability. For example, in non-primary ACs, the majority of neurons could have clear broader tuning properties, which might have reduced the SNR and thus increased the variability of tonotopy mapping results across subjects. Neurons in different parts of the AC might also be sensitive to differing stimulation and task parameters, making the conventional tonotopy mapping less sensitive for mapping subarea boundaries of the higher areas (Phillips et al., 1994). The present results, which suggest that the intrinsic functional connectivity is less variable in HG than in the lateral superior temporal cortex, support the interpretation that these previous observations could not be simply explained by SNR issues.

### The human auditory cortex is not just a simple feature processing area

An intriguing finding of the present study is that, although basic perceptual processes and their cortical substrates vary in the visual domain as well (Farkas et al., 2018), human ACs are significantly more variable across individuals than comparable hierarchical levels of VC. This is consistent with distinct pieces of evidence from previous studies, which have suggested that ACs reflect a higher processing stage, which could be assumed to be more prone to developmental and environmental influences than the corresponding levels of VC processing. For example, there is evidence that certain parts of ACs demonstrate a larger degree of distant vs. local connectivity (Mueller et al., 2013) and are preceded by a larger number of pre-cortical processing steps than comparable cortical stages of visual processing (King and Nelken, 2009; Masterton, 1992). Moreover, it has been suggested that ACs contain neurons with more multidimensional activation preferences for both simple and complex stimulus attributes (Chambers et al., 2014), and encode complex object representations even in the primary input areas (Nelken, 2004). These notions have inspired a theoretical assumption that early human ACs could constitute a higher-level processing center than early visual (or somatosensory) cortices (Nelken et al., 2003), where functional properties may be related to individual variability in auditory-behavioral skills that are uniquely advanced in primates, most prominently so in humans. However, until now, few previous studies had been able to directly compare the degree to which the functional anatomy varies across AC vs. VC areas. A major challenge has been that the properties of the human AC must be characterized using techniques suitable for individual-level studies of dynamic functional networks that also encompass higher cortical areas, rather than using fMRI localizer designs utilized in traditional sensory-cortex mapping at the group level. The present results, which are based on model-free functional connectivity analyses, thus significantly extend current knowledge of the variability of auditory functions as compared to VC functions.

### Functional laterality in human and macaque ACs

In the present study, we observed significantly greater individual variability in the left than right hemisphere in both primate species. This finding is consistent with the presumed hemisphere lateralization of auditory-verbal communication processes in humans, as well as with the relative expansion of left vs. right superior temporal areas that is most prominent in humans (Geschwind and Levitsky, 1968) but also clearly present in simians. Our finding is also in line with results obtained in task-based studies using language-related tasks, which have documented very large individual variability in the activation foci of the left AC areas, comparable to that in the frontal cortices (but see (Bonte et al., 2013) for different interpretations). Further, previous studies suggest that the individual variability of left AC function is correlated with idiosyncrasies of not only fundamental “perceptual styles” (Farkas et al., 2018), but also in voice (Postma-Nilsenová and Postma, 2013) and speech production processes (Franken et al., 2017). Here we showed that neural connectivity at rest is already more variable in the left AC compared to the right AC. This indicates that the unique wiring pattern in the left AC of each subject may be particularly important for understanding individual differences in auditory functions. Given that functional connectivity measured at rest can be related to individual differences in complex cognitive abilities (Finn et al., 2015), we speculate that connectivity in the left AC may provide valuable predictors of speech and language development in both abnormal and normal populations.

### Interspecies comparisons

Previous architectonic studies suggest that although largely homologous AC subregions are found in all primates, the degree of individual variability and complexity are larger in great apes and humans than in monkeys (Hackett et al., 2001). It is thus tempting to conjecture that the human ACs are uniquely complex, also in terms of the patterns of their individual variability. This speculation receives indirect support from surface-based MRI mapping studies quantifying the expansion of different neocortical areas between monkeys and humans (Van Essen and Glasser, 2014), which show indices of larger interspecies expansion of certain higher AC than VC areas. It is also noteworthy that, even when the account of language evolution is disregarded, humans differ from other primates more prominently auditory than visual cognitive skills such as working memory (Scott et al., 2012). However, our data provided strong evidence that the degree of functional variability of monkey brain function, similarly to humans, is greater in the AC than VC and, importantly, is also greater in the left than right AC. The existence of such a human-like cortical distributions of individual variability in our close ancestors could reflect the neurobiological substrate for processing of complex auditory signals that has, ultimately, contributed to the evolution of speech communication in humans.

### Limitations and caveats

Several limitations of the study are worth mentioning. First, functional variability could be confounded by errors in image registration in areas whose folding patterns vary across subjects, such as in HG (for a review, see Moerel et al., 2014). Specifically, a higher degree of convolution can lead to lower fidelity of inter-subject alignment (Van Essen, 2005). To investigate this potential confound, we regressed out sulcal depth variability, which comprises variability due to alignment error, from the functional variability map. We found that the overall pattern of functional connectivity variability remained stable after regression. It is also important to note that the hierarchical organization of inter-subject variability within ACs themselves would likely have looked like something completely different if the fcMRI variability would be a byproduct of anatomical variability only. That is, in contrast to HG whose folding patterns vary substantially across subjects (for a review, see Moerel et al., 2014), STG has been considered structurally quite similar across individuals (Coalson et al., 2018), which makes it relatively robust to align an individual subject’s STG to a standard template. However, the degree of fcMRI inter-subject variability was, nonetheless, much greater at the crest of STG than in HG. Previous studies have also show that surface-based anatomical alignment does a really good job in early VCs (Hinds et al., 2009), whose folding patterns and anatomical size do, in fact, vary at least equally to those of human superior temporal plane. (The fact that the same has not yet been shown in ACs could be explained by the lack of a two-dimensional functional marker of subarea boundaries.) Most importantly, the results found in the human brain were highly consistent to those found in the macaque who do not have a HG.

The second potential limitation of study is that our quantification of variability in task activations is limited to a single dataset that used vocal and non-vocal stimuli; thus, our finding may not generalize to other tasks. Future work based on different auditory tasks is warranted. Third, it also must be noted that human AC subareas might be smaller than those in the VCs, which could have affected the comparison of inter-subject variability between the AC and the VC. Further studies with higher-resolution fMRI techniques that allow for smaller voxel sizes (e.g., sub-millimeter BOLD image using 7 Tesla MRI) could help resolve this issue. Fourth, inter-subject variability in the macaque brain was estimated using the data of only four subjects, although each subject had significant amount of data. To examine whether inter-subject variability is dependent on a large sample size, we randomly selected four human subjects and re-estimated inter-subject variability. We found that the results from four subjects were already quite similar to the results derived from 30 subjects (Pearson correlation r =0.79, see Figure S4). Thus, the data from four monkeys may be able to accurately reflect inter-subject variability. Fifth, inter-subject variability may be affected by different states of consciousness (awake vs. anesthetized) (Xu et al., 2019). Further explorations are needed to gain a better understanding about how the functional variability in auditory cortex is related to consciousness. Finally, to limit the possible impact of acoustical scanner noise on AC functional connectivity, we estimated intra-subject variability based on repeated scans and used it as a regressor. However, it is possible that the noise effect on inter-subject variability is not fully captured by intra-subject variability. One way to test this in future studies could be the sparse temporal sampling technique (Hall et al., 1999). Nevertheless, recent studies using conventional fMRI resolutions show that topographically organized resting-state functional connectivity patterns also emerge in human ACs (Cha et al., 2016). The fact that this arrangement is evident even in congenitally deaf individuals during resting-state fMRI (Striem-Amit et al., 2016) suggests that the result is not explainable by the fluctuations caused by the acoustical noise of fMRI. If anything, the constant background acoustic stimulation should increase the consistency of activation patterns in ACs, as compared to VCs.

## Materials and Methods

### Participants and data collection

Three fMRI datasets obtained with different imaging parameters were employed in the present study.

#### CoRR-HNU dataset

The Hangzhou Normal University of the Consortium for Reliability and Reproducibility (CoRR-HNU) dataset (Zuo et al., 2014) consisted of 30 young healthy adults (15 females, mean age = 24, SD = 2.41). None of the participants had a history of neurological or psychiatric disorders, substance abuse, or head injury with loss of consciousness. Each subject underwent ten 10-min scanning sessions over approximately one month. The ethics committee of the Center for Cognition and Brain Disorders (CCBD) at Hangzhou Normal University approved the study. Written informed consent was obtained from each participant prior to data collection. MRI data were acquired on a GE MR750 3 T scanner (GE Medical Systems, Waukesha, WI, USA). Structural images were acquired using a T1-weighted Fast Spoiled Gradient echo (FSPGR: TR = 8.1 ms, TE = 3.1 ms, TI = 450 ms, flip angle = 8°, field of view = 256 × 256 mm, matrix = 256 × 256, voxel size = 1.0 × 1.0 × 1.0 mm, 176 sagittal slices). Functional data were obtained using an echo-planar imaging sequence (EPI: TR = 2000 ms, TE = 30 ms, flip angle = 90°, field of view = 220 × 220 mm, matrix = 64 × 64, voxel size = 3.4 × 3.4 × 3.4 mm, 43 slices). The participants were instructed to relax and remain still with their eyes open, not to fall asleep, and not to think about anything in particular. The screen presented a black crosshair in the center of a gray background.

#### MSC dataset

The Midnight Scanning Club (MSC) dataset (Gordon et al., 2017) included 10 healthy young adults (5 females, mean age = 29.1, SD = 3.3). Informed consent was obtained from all participants. The study was approved by the Washington University School of Medicine Human Studies Committee and Institutional Review Board. For each participant, 30 continuous minutes of resting state were scanned on 10 separate days on a Siemens TRIO 3 T MRI scanner (Erlangen, Germany). Structural MRI data was obtained using T1-weighted images (voxel size = 1.0 × 1.0 × 1.0 mm, TE = 3.74 ms, TR = 2400 ms, TI = 1000 ms, flip angle = 8°, 224 sagittal slices). All functional imaging data was acquired using a gradient-echo EPI sequence (TR = 2.2 s, TE = 27 ms, flip angle = 90°, voxel size = 4mm × 4mm × 4 mm, 36 slices). The participants visually fixated on a white crosshair presented against a black background.

#### Human-voice dataset

The task dataset (Pernet et al., 2015) included 218 healthy adults (117 males; mean age = 24.1, SD = 7.0). Participants all provided written informed consent prior to participation, in accordance with the Declaration of Helsinki. The experiments were approved by the local ethics committee at the University of Glasgow. All fMRI data were acquired from a Siemens TRIO 3 T MRI scanner (Erlangen, Germany) using a single-shot gradient-echo echo-planar imaging sequence (EPI, TR = 2000 ms, TE = 30 ms, flip angle = 77°, field of view = 210 × 210 mm, matrix = 70 × 70, voxel size = 3 × 3 × 3.3 mm, 32 slices). In addition to the 310 EPI volumes, a high-resolution 3D T1-weighted sagittal scan was obtained for each subject (voxel size = 1.0 × 1.0 × 1.0 mm, matrix = 256 × 256 × 192). Each run consisted of 10 min and 20 s block design with forty 8-s long blocks of either vocal (20 blocks) or non-vocal (20 blocks) sounds. The vocal or non-vocal blocks were intermixed randomly with 20 periods of silence. Subjects were scanned while passively listening to the stimuli and keeping their eyes closed. Other details of the data collection and task design can be found elsewhere (Pernet et al., 2015).

#### Macaque dataset I & II

Macaque dataset I included two rhesus monkeys (Macaca mulatta, one male, age 6 years, 6.4 kg; one female, age 7 years, 4.5 kg), which was collected from the Nathan Kline Institute for Psychiatric Research. All methods and procedures were approved by the NKI Institutional Animal Care and Use Committee (IACUC) protocol. MRI images were acquired using a Simens Tim Trio 3T MRI scanner with an 8-channel surface coil adapted for the monkeys’ head. Structural MRI images were acquired using T1-weighted images (0.5 mm isotropic voxel, TE = 3.87 ms, TR = 2500 ms, TI=1200 ms, flip angle=8 degrees). All functional images were acquired utilizing a gradient echo EPI sequence (TR=2000 ms, TE=16.6 ms, flip angle = 45 degree, 1.5 × 1.5 × 2mm voxels, 32 slices, FOV = 96 × 96 mm). For each macaque, eight resting-state scans (10 min for each scan) from 2 anesthetized sessions were collected with Monocrystalline iron oxide ferumoxytol (MION).

Macaque dataset II included two male rhesus macaques (Macaca mulatta, one male, age 5 years, 8.6 kg; one male, age 5 years, 7.6 kg), which was collected from the Oregon Health and Science University. Animal procedures were in accordance with the National Institutes of Health guidelines on the ethical use of animals and were approved by the Oregon National Primate Research Center (ONPRC) Institutional Animal Care and Use Committee. MRI images were acquired using a Simens Tim Trio 3T MRI scanner with a 15-channel coil adapted for the monkeys’ head. Structural MRI images were obtained using T1-weighted images (0.5 mm isotropic voxel, TE = 3.33 ms, TR = 2600 ms, TI = 900 ms, flip angle = 8 degrees). All functional data were acquired using a gradient echo EPI sequence (TR = 2070 ms, TE = 25 ms, flip angle = 90 degrees, 1.5 × 1.5 × 1.5 mm voxels, 32 slices, FOV = 96 × 96 mm). For each macaque, eight 30-min anesthetized scans were acquired with MION.

Other details of the data collection can be found in previous reports of the datasets (Xu et al., 2018).

### Data Preprocessing

#### CoRR-HNU dataset

Resting-state fMRI data of the 30 subjects in this dataset were processed using the procedures previously described (Mueller et al., 2013). The following steps were performed: (i) slice timing correction (SPM2; Wellcome Department of Cognitive Neurology, London, UK), (ii) rigid body correction for head motion with the FSL package, (iii) normalization for global mean signal intensity across runs, and (iv) band-pass temporal filtering (0.01–0.08 Hz), head-motion regression, whole-brain signal regression, and ventricular and white-matter signal regression.

Structural data were processed using FreeSurfer version 5.3.0. Surface mesh representations of the cortex from each individual subject’s structural images were reconstructed and registered to a common spherical coordinate system. The structural and functional images were aligned using boundary-based registration within the FsFast software package (http://surfer.nmr.mgh.harvard.edu/fswiki/FsFast). The preprocessed resting-state BOLD fMRI data were then aligned to the common spherical coordinate system via sampling from the middle of the cortical ribbon in a single interpolation step. FMRI data of each individual were registered to the FreeSurfer cortical surface template (fsaverage6) that consists of 40,962 vertices in each hemisphere. A 6-mm full-width half-maximum (FWHM) smoothing kernel was then applied to the fMRI data in the surface space.

#### MSC dataset

Resting-state fMRI data and structural data of the 10 subjects in this dataset were preprocessed identically to the *CoRR-HNU dataset*.

#### Human-voice dataset

Conventional task-evoked activation maps in this dataset were estimated using FSL’s FEAT (https://fsl.fmrib.ox.ac.uk/fsl/fslwiki/FEAT). After slice timing, rigid body correction and high-pass temporal filtering (100 Hz), task-induced BOLD responses were modeled by convolving the double-gamma hemodynamic response function with the experimental design. Structural data of the 218 subjects in this dataset were preprocessed identically to the *CoRR-HNU dataset*. The task-evoked activation maps of each individual were also projected to fsaverage6.

#### Macaque dataset

The procedure of the structural data was similar with that of the human datasets but was edited manually during the tissue segmentation and the surface reconstruction. After generating the native white-matter and pial surfaces by using FreeSurfer, we then registered the native surfaces to a hybrid left-right template surface (Yerkes19 macaque template (Donahue et al., 2016)).

Resting-state fMRI data were processed by slice timing correction, motion correction and bias filed correction (for *Macaque dataset II*), band-pass temporal filtering (0.01-0.1 Hz). Head-motion parameters, white-matter, ventricular and whole-brain signals were linearly regressed out. We then transformed the denoised functional images into the corresponding anatomical images and then into the native mid-thickness surface. A 4 mm FWHM smoothing kernel was then applied on the native surface. The smoothed data were downsampled to the 10k (10,242 vertices) Yerkes19 template surface. More details about the preprocessing procedure of the macaque datasets can be found in the previous report (Xu et al., 2018).

### Generating Masks for Auditory and Visual Cortices

For the human data, the auditory cortex mask was described in our previous paper (Ahveninen et al., 2016) and the visual cortex mask was from the published V1-V3 visual cortex mask (Benson et al., 2014). The Left auditory, right auditory and left visual masks included 2,155 vertices, 1,984 vertices and 2,810 vertices, respectively. The cortical surface was downsampled to 1,175 vertices (i.e., regions of interests, ROIs) that were approximately uniformly distributed across the two hemispheres.

For the macaque data, the auditory cortex and visual cortex masks were extracted from the Markov’s cytoarchitectonic cortical parcellation (see Figure S3) (Markov et al., 2012). The auditory cortex consisted of Core, Lateral Belt (LB), Medial Belt (MB), caudal-part and rostral-part Parabelt (PBc and PBr) areas. The visual cortex consisted of V1, V2 and V3 parcels. The left auditory, right auditory and left visual masks included 266 vertices, 246 vertices, 1,684 vertices, respectively. The cortical surface was downsampled to 1,112 uniformly distributed ROIs, which were generated using similar methods as for the human data.

#### Estimating Inter-Individual Variability of Resting-state Functional Connectivity within the Auditory and Visual Cortices

BOLD fMRI signal time courses were extracted from the auditory and visual cortex masks, respectively. Functional connectivity profiles were obtained by computing Pearson’s correlation between time courses of the vertices within each mask and time courses of the cortical ROIs. The profile for a given vertex *i* could be denoted as *F*_*i*_ (*s, v*), where *i* = 1,2…*N*, and *F*_*i*_ is a 1 × 1175 (or 1112 for macaques) vector, *s* indicates the subject, *v* indicates the session and *N* indicates the number of vertices within the masks. For a given vertex *i*, the intrasubject variance was estimated using the *V* maps derived from all *V* sessions of each subjects (e.g., V=10 for both CoRR-HNU and MSC data):

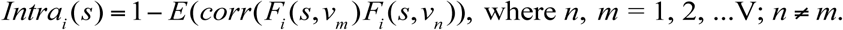

The intrasubject variance was then averaged across all subjects within any one dataset:

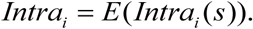

The similarity between *S* (the number of subjects within each dataset, e.g. *S* = 30 in the *CoRR-HNU dataset* while *S* = 10 in the *MSC dataset*) maps derived from all subjects was quantified by averaging the correlation maps between any two maps:

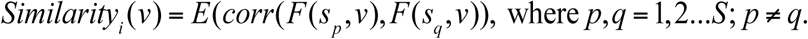

To estimate inter-individual variability, the similarity map was inverted (by subtraction from 1) and then the intrasubject variance was regressed out using a general linear model (GLM). The residual map could be regarded as the inter-individual variability of resting-state functional connectivity:

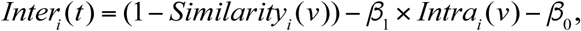

where *β*_1_ and a *β*_0_ are parameters determined by the GLM. Inter-individual variability maps derived from each session *t* are averaged.

#### Estimating Inter-individual Variability of Task Activation within the Auditory Cortex

The task activation *z*-value maps derived from the *Human-voice dataset* were extracted from the same auditory cortex mask that was used for resting-state functional connectivity variability. The standard deviation of the *z* values for each task contrast across all 218 subjects was calculated to estimate inter-individual variance, while the average *z*-value map was estimated by averaging task activation *z*-value maps across all subjects. Normalization (*z*-score) was then applied to both the standard deviation map and the mean *z*-value map derived from all subjects within the auditory cortex mask. The normalized mean *z*-value map was regressed out from the normalized inter-individual standard deviation map to wean off its dependence on the mean *z*-value. The resulting residual map may be considered the inter-individual variability of the task fMRI data.

#### Relationship to Anatomical Variability

Sulcal depth and cortical thickness measurements were calculated using FreeSurfer. The sulcal depth estimated by FreeSurfer is the integrated dot product of the movement vector with the surface normal during inflation. It highlights large-scale geometry as deep regions consistently move outward and have a positive value while superficial regions move inward and have a negative value. Inter-individual variability in sulcal depth and cortical thickness was estimated vertex-wise using intraclass correlation (ICC) with the intrasubject variance regression. Pearson’s correlation coefficient was calculated between functional variability and anatomical variability across the auditory cortex.

#### Inter-subject variability in Seed-based Functional Connectivity

In order to visualize the differences of the functional connectivity patterns between seed in the high-variability region and seed in the low-variability region, we selected two juxtaposed seeds in the AC but one of them located in the low-variability region around HG (MNI coordinate: −60, −18, 1) and another located in the high-variability region in STG (MNI coordinate: −62, −18, −2). We estimated the seed-based functional connectivity maps for every single individual by using Pearson’s product moment correlation. We then converted them to *z*-maps using Fisher’s r-to-z transformation and averaged the *z*-maps across all 30 subjects (Figure S1).

#### Statistics

Wilcoxon rank sum tests were used to compare the functional variability between the AC and the VC, and between the left and right ACs. Pearson’s correlations were used to evaluate the relationship between variability in functional connectivity and variability in anatomical features. To test the potential impact of spatial dependence between neighboring vertices on correlation analysis, we performed a repeated (n = 1,000) random sampling of 7% of the vertices and computed the correlation coefficient on the subsets of the vertices. For each subset, the Durbin-Watson test was performed to estimate the spatial dependence (DW > 2). Correlation coefficients were averaged across the 1,000 iterations.

#### Visualization

All results were projected on the Freesurfer cortical surface template “fsaverage” for visualization purposes. The VC was cut along the calcarine fissure and flattened using the FreeSurfer command (mris_flatten).

## Data and code availability

The CoRR-HNU dataset is publically available through Consortium for Reliability and Reproducibility Project (http://fcon_1000.projects.nitrc.org/indi/CoRR/html/hnu_1.html). The MSC dataset is publicly available through OpenfMRI (https://openfmri.org/dataset/ds000224/). The Human-voice dataset is also publicly available through OpenfMRI (https://openfmri.org/dataset/ds000158/). MATLAB codes that support the findings of this study are available from the corresponding authors upon request.

## Acknowledgements

We would like to thank the investigative teams from the Nathan Kline Institute (C. Schroeder, M.P. Milham, A. Falchier, S. Colcombe, G. Linn, D. Ross, R.C. Craddock), Oregon Health and Science University (E.L. Sullivan, E. Feczko, J. Bagley, E. Earl, O. Miranda-Domingue, D. Fair), as well as the funding agencies that made their work possible (NKI: NIH; OHSU: NIH and NIMH).

## Funding

This work was supported by NIH grants R01NS091604, P50MH106435, R01DC016765, and R01DC016915; and by the National Natural Science Foundation of China grant No. 81790652, 81522021 and 81671662.

## Author contributions

All authors contributed to the conception and design of the work, as well as the analysis and interpretation of the data.

## Data and materials availability

All data needed to evaluate the conclusions in the paper are present in the paper and/or the Supplementary Materials. All data related to this paper may be requested from the authors.

## Competing interests

The authors declare that they have no competing interests.

## Supplemental Figures

**Figure S1.**
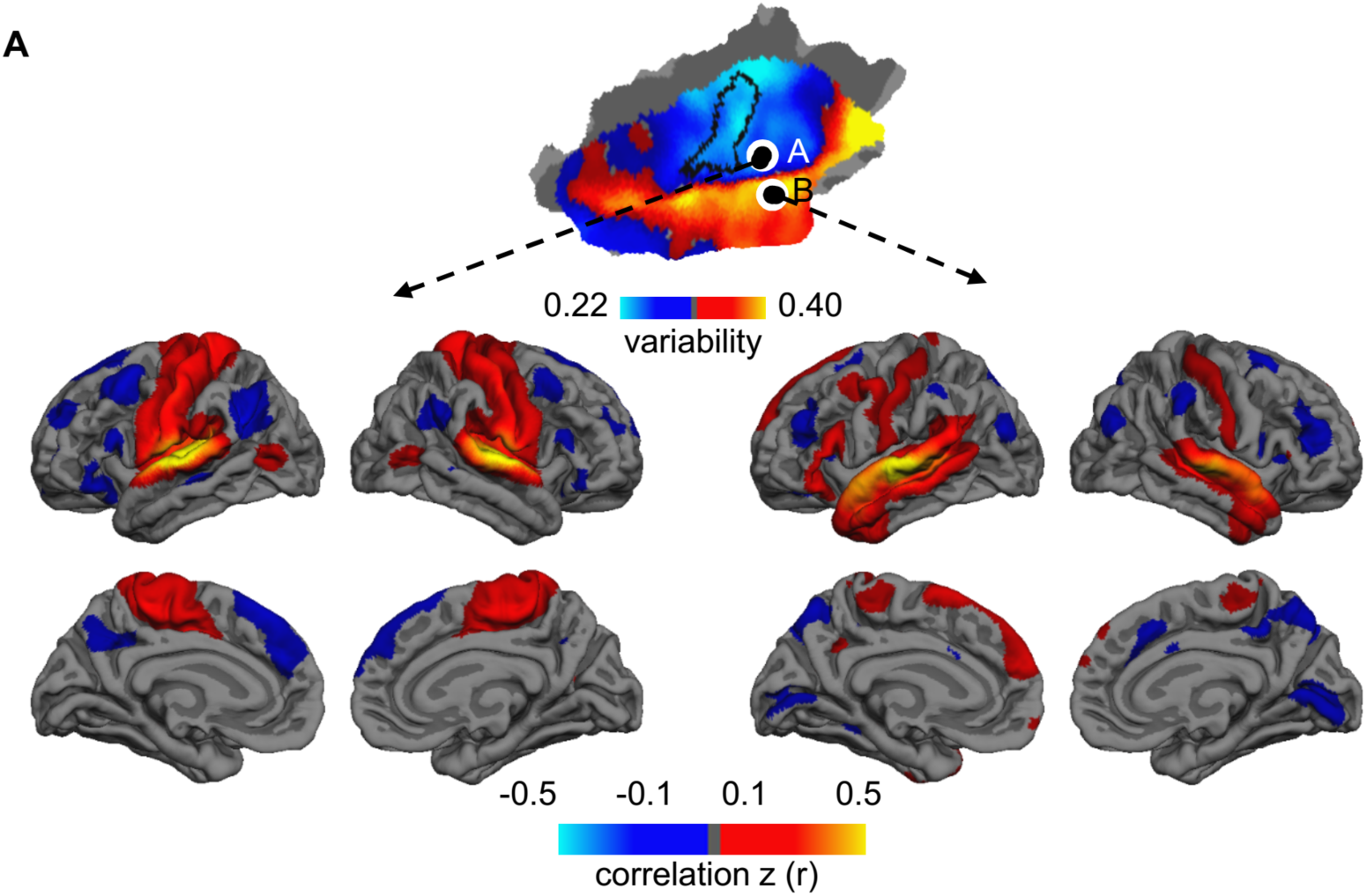
Related to Figure 1. Auditory regions with low variability and high variability show distinct functional connectivity patterns. Two seeds were placed in the human AC, one in the regions showing low inter-individual variability (seed A) and the other in the region showing high variability (seed B). Group-level functional connectivity maps were derived using these two seeds. Although the two seeds were very close to each other, the seed in the low variability region is strongly connected to the sensorimotor cortex, whereas the seed in the high variability region shows strong connectivity to the inferior frontal gyrus and temporal pole. These distinct connectivity patterns suggest that auditory regions with low variability might involve the primary information processing whereas regions with high variability might involve higher-order association processing including language functions.

**Figure S2.**
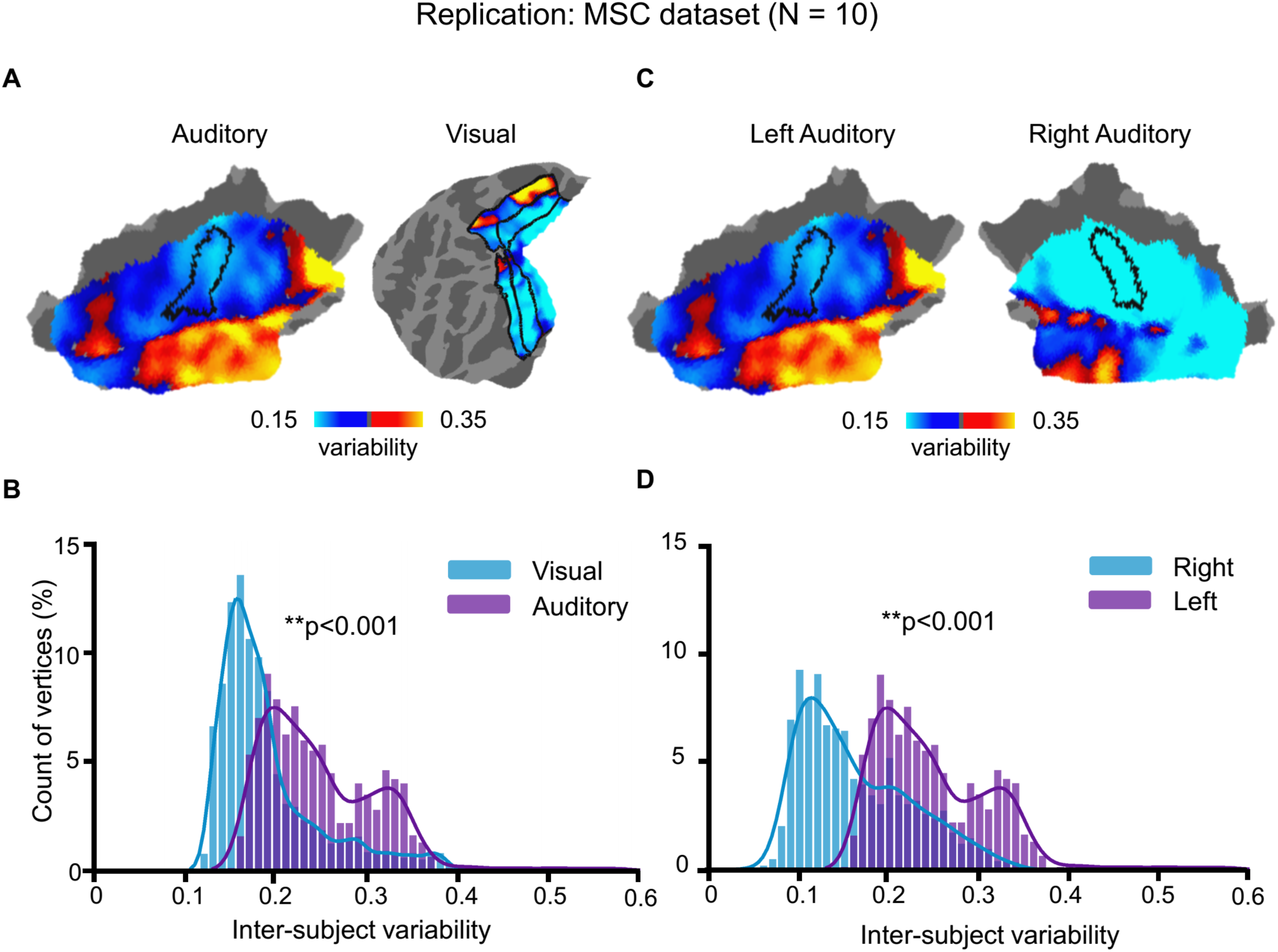
Related to Figure 1 and Figure 2. The greater variability of auditory than visual cortex and left lateralization in the auditory cortex were replicated in an independent dataset. The main findings derived from the CoRR-HNU dataset (N = 30) were replicated in an independent human dataset (MSC Dataset, N=10) with different scanning parameters and subjects’ ethnicities. Inter-individual variability of the AC derived from the two datasets was highly similar (Pearson correlation r = 0.836, p <0.0001). Furthermore, we replicated the findings that **(A, B)** variability in the AC is greater than that in the VC (p < 0.001, Wilcoxon Rank Sum test) and **(C, D)** variability in the left AC is significantly greater than that in the right AC (p < 0.001, Wilcoxon Rank Sum test).

**Figure S3.**
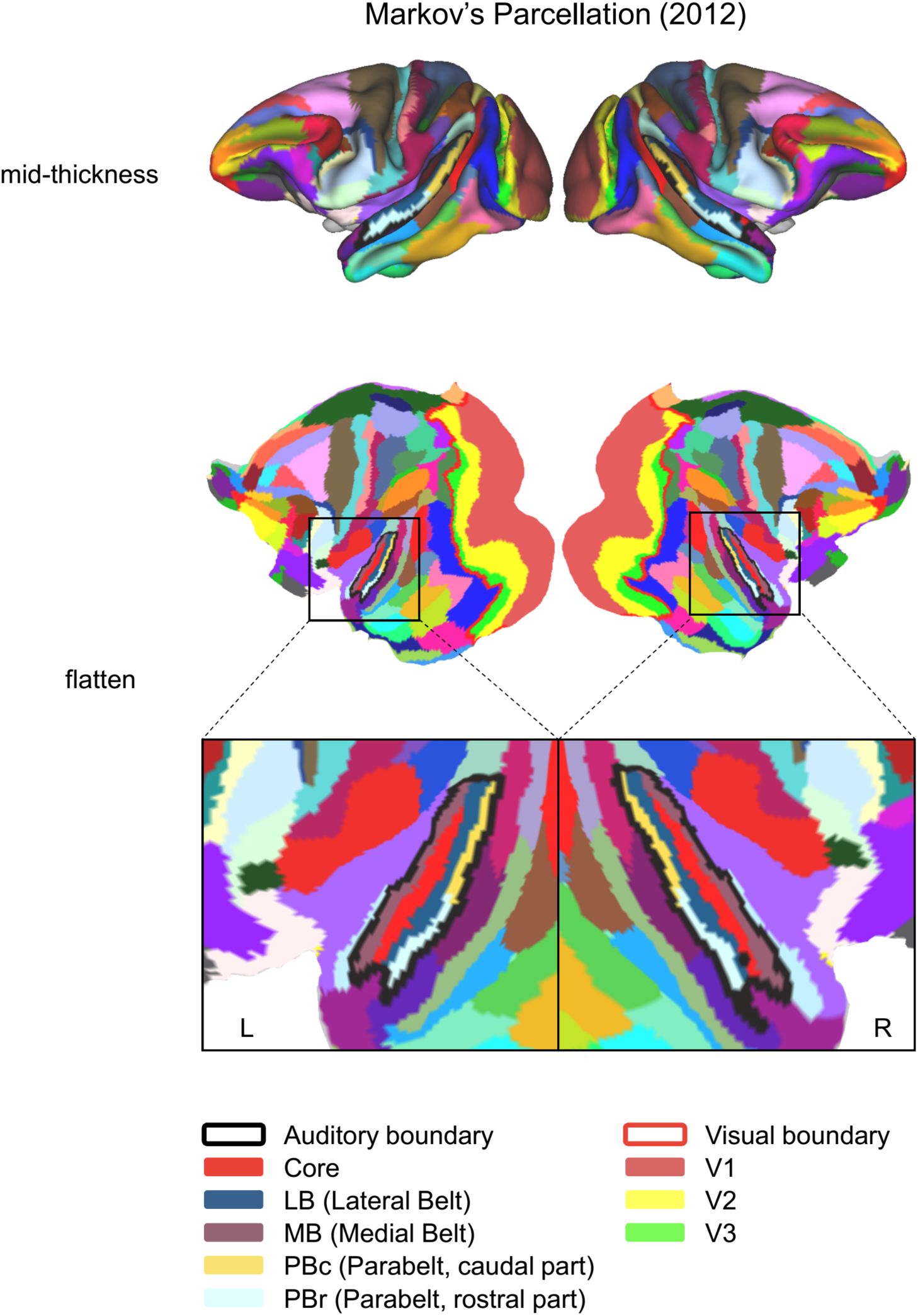
Related to Figure 1 and Figure 2. Markov’s cytoarchitectonie cortical parcellation was used to generate the masks of the auditory and visual cortices in macaques. Markov’s cortical parcellation is shown on the mid-thickness surface (upper panel) and flatten surfaces (middle panel) of both hemispheres. A red curve delineates the boundary of the visual cortex, consisting of V1-V3 areas. A black curve delineates the boundary of the auditory cortex, consisting of the core, lateral and medial belt, caudal and rostral parabelt areas. The auditory cortex is magnified to demonstrate more details (bottom panel).

**Figure S4.**
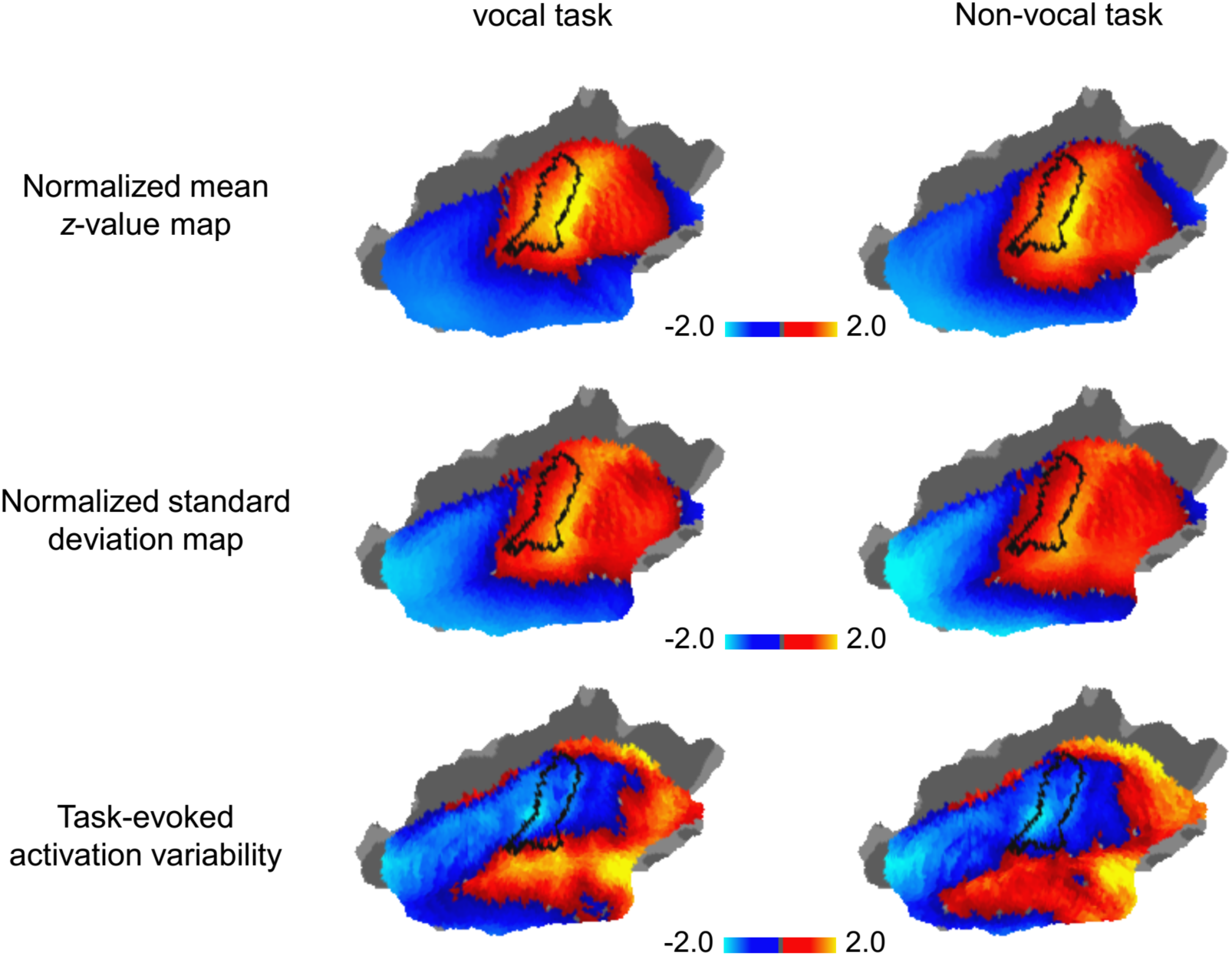
Related to Figure 3. Mean, standard deviation, and variability of task-evoked activations were evaluated using the vocal and non-vocal tasks. The normalized average *z*-value maps derived from both vocal stimulus (left column) and non-vocal stimulus (right column) represent the baseline of task-evoked activations (upper panel). Standard deviation maps represent the raw inter-individual variability of the task-evoked activations without accounting for the baseline distribution (middle panel). Inter-individual variability of the task-evoked activations was evaluated by linearly regressing out the baseline from the standard deviation maps (bottom panel).

**Figure S5.**
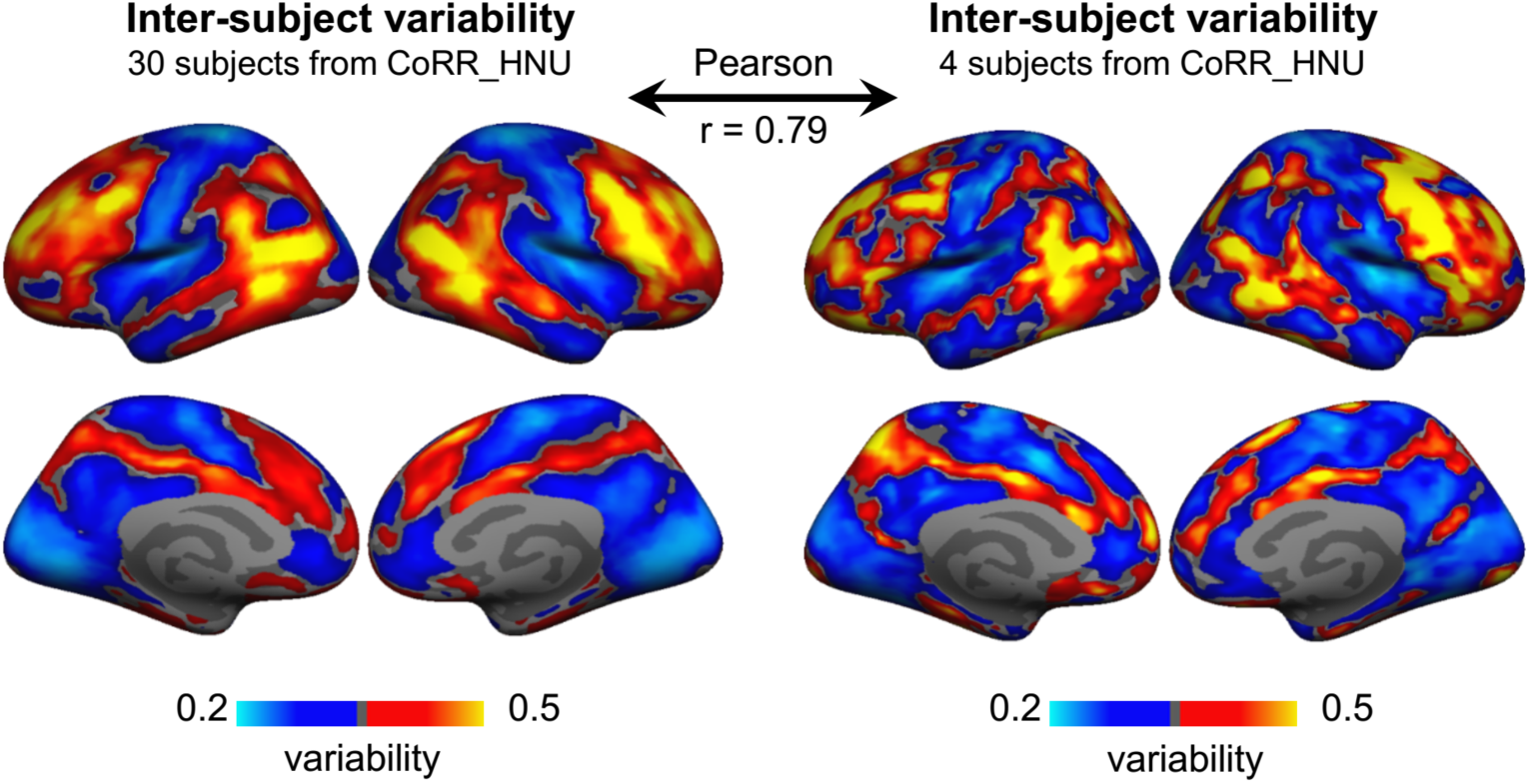
Inter-individual variability can be robustly estimated using a small sample. Due to the limited sample size of the macaque dataset (N = 4), it is necessary to investigate whether a small sample can be used to estimate individual variability in functional connectivity. Four subjects were randomly picked up from the CoRR-HNU dataset to evaluate individual variability. Variability derived from four subjects (right column) is highly similar to that derived from 30 subjects (left column, Pearson correlation r = 0.79, p < 0.0001), suggesting that inter-individual variability can be robustly estimated using a small sample.

## REFERENCES

Aboitiz, F. (2018). A Brain for Speech. Evolutionary Continuity in Primate and Human Auditory-Vocal Processing. Front Neurosci 12, 174.

Ahveninen, J., Chang, W.-T., Huang, S., Keil, B., Kopco, N., Rossi, S., Bonmassar, G., Witzel, T., and Polimeni, J.R. (2016). Intracortical depth analyses of frequency-sensitive regions of human auditory cortex using 7T fMRI. Neuroimage 143, 116–127.

Arcadi, A.C. (1996). Phrase structure of wild chimpanzee pant hoots: Patterns of production and interpopulation variability. American Journal of Primatology 39, 159–178.

Belin, P. (2006). Voice processing in human and non-human primates. Philos Trans R Soc Lond B Biol Sci 361, 2091–2107.

Benson, N.C., Butt, O.H., Brainard, D.H., and Aguirre, G.K. (2014). Correction of distortion in flattened representations of the cortical surface allows prediction of V1-V3 functional organization from anatomy. PLoS computational biology 10, e1003538.

Bizley, J.K., and Cohen, Y.E. (2013). The what, where and how of auditory-object perception. Nat Rev Neurosci 14, 693–707.

Bonte, M., Frost, M.A., Rutten, S., Ley, A., Formisano, E., and Goebel, R. (2013). Development from childhood to adulthood increases morphological and functional inter-individual variability in the right superior temporal cortex. Neuroimage 83, 739–750.

Cha, K., Zatorre, R.J., and Schonwiesner, M. (2016). Frequency Selectivity of Voxel-by-Voxel Functional Connectivity in Human Auditory Cortex. Cereb Cortex 26, 211–224.

Chambers, A.R., Hancock, K.E., Sen, K., and Polley, D.B. (2014). Online stimulus optimization rapidly reveals multidimensional selectivity in auditory cortical neurons. J Neurosci 34, 8963–8975.

Cheung, S.W., Nagarajan, S.S., Schreiner, C.E., Bedenbaugh, P.H., and Wong, A. (2005). Plasticity in primary auditory cortex of monkeys with altered vocal production. J Neurosci 25, 2490–2503.

Coalson, T.S., Van Essen, D.C., and Glasser, M.F. (2018). The impact of traditional neuroimaging methods on the spatial localization of cortical areas. Proc Natl Acad Sci U S A 115, E6356–E6365.

De Martino, F., Moerel, M., Xu, J., van de Moortele, P.F., Ugurbil, K., Goebel, R., Yacoub, E., and Formisano, E. (2015). High-Resolution Mapping of Myeloarchitecture In Vivo: Localization of Auditory Areas in the Human Brain. Cereb Cortex 25, 3394–3405.

Dick, F., Tierney, A.T., Lutti, A., Josephs, O., Sereno, M.I., and Weiskopf, N. (2012). In vivo functional and myeloarchitectonic mapping of human primary auditory areas. J Neurosci 32, 16095–16105.

Donahue, C.J., Sotiropoulos, S.N., Jbabdi, S., Hernandez-Fernandez, M., Behrens, T.E., Dyrby, T.B., Coalson, T., Kennedy, H., Knoblauch, K., and Van Essen, D.C. (2016). Using diffusion tractography to predict cortical connection strength and distance: a quantitative comparison with tracers in the monkey. Journal of Neuroscience 36, 6758–6770.

Farkas, D., Denham, S.L., and Winkler, I. (2018). Functional brain networks underlying idiosyncratic switching patterns in multi-stable auditory perception. Neuropsychologia 108, 82–91.

Finn, E.S., Shen, X., Scheinost, D., Rosenberg, M.D., Huang, J., Chun, M.M., Papademetris, X., and Constable, R.T. (2015). Functional connectome fingerprinting: identifying individuals using patterns of brain connectivity. Nature neuroscience 18, 1664.

Fischl, B., and Sereno, M.I. (2018). Microstructural parcellation of the human brain. Neuroimage 182, 219–231.

Franken, M.K., Acheson, D.J., McQueen, J.M., Eisner, F., and Hagoort, P. (2017). Individual variability as a window on production-perception interactions in speech motor control. J Acoust Soc Am 142, 2007.

Geschwind, N., and Levitsky, W. (1968). Human brain: left-right asymmetries in temporal speech region. Science 161, 186–187.

Ghazanfar, A.A., and Santos, L.R. (2003). Primates as auditory specialists. In Primate audition Ethology and neurobiology, A.A. Ghazanfar, ed. (Boca Raton, FL: CRC Press), pp. 1–12.

Gordon, E.M., Laumann, T.O., Gilmore, A.W., Newbold, D.J., Greene, D.J., Berg, J.J., Ortega, M., Hoyt-Drazen, C., Gratton, C., and Sun, H. (2017). Precision functional mapping of individual human brains. Neuron 95, 791-807. e797.

Griffiths, T.D., and Warren, J.D. (2002). The planum temporale as a computational hub. Trends in Neurosciences 25, 348–353.

Hackett, T.A., Preuss, T.M., and Kaas, J.H. (2001). Architectonic identification of the core region in auditory cortex of macaques, chimpanzees, and humans. Journal of Comparative Neurology 441, 197–222.

Hall, D.A., Haggard, M.P., Akeroyd, M.A., Palmer, A.R., Summerfield, A.Q., Elliott, M.R., Gurney, E.M., and Bowtell, R.W. (1999). “Sparse” temporal sampling in auditory fMRI. Human brain mapping 7, 213–223.

Herholz, S.C., and Zatorre, R.J. (2012). Musical training as a framework for brain plasticity: behavior, function, and structure. Neuron 76, 486–502.

Hill, J., Dierker, D., Neil, J., Inder, T., Knutsen, A., Harwell, J., Coalson, T., and Van Essen, D. (2010). A surface-based analysis of hemispheric asymmetries and folding of cerebral cortex in term-born human infants. J Neurosci 30, 2268–2276.

Hinds, O., Polimeni, J.R., Rajendran, N., Balasubramanian, M., Amunts, K., Zilles, K., Schwartz, E.L., Fischl, B., and Triantafyllou, C. (2009). Locating the functional and anatomical boundaries of human primary visual cortex. Neuroimage 46, 915–922.

Kaas, J.H. (2006). Evolution of the neocortex. Curr Biol 16, R910–914.

Kell, A.J.E., and McDermott, J.H. (2019). Invariance to background noise as a signature of non-primary auditory cortex. Nat Commun 10, 3958.

Kenet, T., Bibitchkov, D., Tsodyks, M., Grinvald, A., and Arieli, A. (2003). Spontaneously emerging cortical representations of visual attributes. Nature 425, 954–956.

King, A.J., and Nelken, I. (2009). Unraveling the principles of auditory cortical processing: can we learn from the visual system? Nat Neurosci 12, 698–701.

Lu, T., and Wang, X. (2004). Information content of auditory cortical responses to time-varying acoustic stimuli. J Neurophysiol 91, 301–313.

Lumaca, M., Kleber, B., Brattico, E., Vuust, P., and Baggio, G. (2019). Functional connectivity in human auditory networks and the origins of variation in the transmission of musical systems. Elife 8.

Markov, N.T., Ercsey-Ravasz, M., Ribeiro Gomes, A., Lamy, C., Magrou, L., Vezoli, J., Misery, P., Falchier, A., Quilodran, R., and Gariel, M. (2012). A weighted and directed interareal connectivity matrix for macaque cerebral cortex. Cerebral cortex 24, 17–36.

Masterton, R.B. (1992). Role of the central auditory system in hearing: the new direction. Trends Neurosci 15, 280–285.

Mesgarani, N., David, S.V., Fritz, J.B., and Shamma, S.A. (2008). Phoneme representation and classification in primary auditory cortex. J Acoust Soc Am 123, 899–909.

Moerel, M., De Martino, F., and Formisano, E. (2014). An anatomical and functional topography of human auditory cortical areas. Front Neurosci 8, 225.

Moerel, M., De Martino, F., Santoro, R., Ugurbil, K., Goebel, R., Yacoub, E., and Formisano, E. (2013). Processing of natural sounds: characterization of multipeak spectral tuning in human auditory cortex. J Neurosci 33, 11888–11898.

Mueller, S., Wang, D., Fox, M.D., Yeo, B.T., Sepulcre, J., Sabuncu, M.R., Shafee, R., Lu, J., and Liu, H. (2013). Individual variability in functional connectivity architecture of the human brain. Neuron 77, 586–595.

Nelken, I. (2004). Processing of complex stimuli and natural scenes in the auditory cortex. Curr Opin Neurobiol 14, 474–480.

Nelken, I., Fishbach, A., Las, L., Ulanovsky, N., and Farkas, D. (2003). Primary auditory cortex of cats: feature detection or something else? Biol Cybern 89, 397–406.

Norman-Haignere, S., Kanwisher, N.G., and McDermott, J.H. (2015). Distinct Cortical Pathways for Music and Speech Revealed by Hypothesis-Free Voxel Decomposition. Neuron 88, 1281–1296.

Pernet, C.R., McAleer, P., Latinus, M., Gorgolewski, K.J., Charest, I., Bestelmeyer, P.E., Watson, R.H., Fleming, D., Crabbe, F., Valdes-Sosa, M., and Belin, P. (2015). The human voice areas: Spatial organization and inter-individual variability in temporal and extra-temporal cortices. Neuroimage 119, 164–174.

Phillips, D.P., Semple, M.N., Calford, M.B., and Kitzes, L.M. (1994). Level-dependent representation of stimulus frequency in cat primary auditory cortex. Exp Brain Res 102, 210–226.

Postma-Nilsenová, M., and Postma, E. (2013). Auditory perception bias in speech imitation. Frontiers in Psychology 4.

Rauschecker, J.P., and Scott, S.K. (2009). Maps and streams in the auditory cortex: nonhuman primates illuminate human speech processing. Nat Neurosci 12, 718–724.

Ressel, V., Pallier, C., Ventura-Campos, N., Diaz, B., Roessler, A., Avila, C., and Sebastian-Galles, N. (2012). An effect of bilingualism on the auditory cortex. J Neurosci 32, 16597–16601.

Salmi, R., Hammerschmidt, K., and Doran-Sheehy, D.M. (2014). Individual distinctiveness in call types of wild western female gorillas. PLoS One 9, e101940.

Scott, B.H., Mishkin, M., and Yin, P. (2012). Monkeys have a limited form of short-term memory in audition. Proc Natl Acad Sci U S A 109, 12237–12241.

Sereno, M., Dale, A., Reppas, J., Kwong, K., Belliveau, J., Brady, T., Rosen, B., and Tootell, R. (1995). Borders of multiple visual areas in humans revealed by functional magnetic resonance imaging. Science 268, 803–804.

Seung, S. (2012). Connectome: How the Brain’s Wiring Makes Us Who We Are (New York: Houghton Mifflin Harcourt Publishing Company).

Stoecklein, S., Hilgendorff, A., Li, M., Förster, K., Flemmer, A.W., Galiè, F., Wunderlich, S., Wang, D., Stein, S., and Ehrhardt, H. (2019). Variable functional connectivity architecture of the preterm human brain: Impact of developmental cortical expansion and maturation. Proceedings of the National Academy of Sciences.

Striem-Amit, E., Almeida, J., Belledonne, M., Chen, Q., Fang, Y., Han, Z., Caramazza, A., and Bi, Y. (2016). Topographical functional connectivity patterns exist in the congenitally, prelingually deaf. Sci Rep 6, 29375.

Tavor, I., Jones, O.P., Mars, R., Smith, S., Behrens, T., and Jbabdi, S. (2016). Task-free MRI predicts individual differences in brain activity during task performance. Science 352, 216–220.

Van Essen, D.C. (2005). A population-average, landmark-and surface-based (PALS) atlas of human cerebral cortex. Neuroimage 28, 635–662.

Van Essen, D.C., and Glasser, M.F. (2014). In vivo architectonics: a cortico-centric perspective. Neuroimage 93 Pt 2, 157–164.

Wang, D., Buckner, R.L., Fox, M.D., Holt, D.J., Holmes, A.J., Stoecklein, S., Langs, G., Pan, R., Qian, T., and Li, K. (2015). Parcellating cortical functional networks in individuals. Nature neuroscience 18, 1853.

Xu, T., Falchier, A., Sullivan, E.L., Linn, G., Ramirez, J.S., Ross, D., Feczko, E., Opitz, A., Bagley, J., and Sturgeon, D. (2018). Delineating the macroscale areal organization of the macaque cortex in vivo. Cell reports 23, 429–441.

Xu, T., Sturgeon, D., Ramirez, J.S., Froudist-Walsh, S., Margulies, D.S., Schroeder, C.E., Fair, D.A., and Milham, M.P. (2019). Inter-individual Variability of Functional Connectivity in Awake and Anesthetized Rhesus Monkeys. Biological Psychiatry: Cognitive Neuroscience and Neuroimaging.

Zuo, X.-N., Anderson, J.S., Bellec, P., Birn, R.M., Biswal, B.B., Blautzik, J., Breitner, J.C., Buckner, R.L., Calhoun, V.D., and Castellanos, F.X. (2014). An open science resource for establishing reliability and reproducibility in functional connectomics. Scientific data 1, 140049.

